# Expression of mini-G proteins specifically halt cognate GPCR trafficking and intracellular signalling

**DOI:** 10.1101/2021.11.24.469908

**Authors:** Yusman Manchanda, Zenouska Ramchunder, Maria M Shchepinova, Guy A Rutter, Asuka Inoue, Edward W Tate, Ben Jones, Alejandra Tomas

## Abstract

Mini-G proteins are engineered thermostable variants of Gα subunits designed to specifically stabilise G protein-coupled receptors (GPCRs) in their active conformation for structural analyses. Due to their smaller size and ease of use, they have become popular tools in recent years to assess specific GPCR behaviours in cells, both as reporters of receptor coupling to each G protein subtype and for in-cell assays designed to quantify compartmentalised receptor signalling from a range of subcellular locations. Here, we describe a previously unappreciated consequence of the co-expression of mini-G proteins with their cognate GPCRs, namely a profound disruption in GPCR trafficking and intracellular signalling caused by the co-expression of the specific mini-G subtype coupled to the affected receptor. We studied the Gαs-coupled pancreatic beta cell class B GPCR glucagon-like peptide-1 receptor (GLP-1R) as a model to describe in detail the molecular consequences derived from this effect, including a complete halt in β-arrestin-2 recruitment and receptor internalisation, despite near-normal levels of receptor GRK2 recruitment and lipid nanodomain segregation, as well as the disruption of endosomal GLP-1R signalling by mini-G_s_ co-expression. We also extend our analysis to a range of other prototypical GPCRs covering the spectrum of Gα subtype coupling preferences, to unveil a widely conserved phenomenon of GPCR internalisation blockage by specific mini-G proteins coupled to a particular receptor. Our results have important implications for the design of methods to assess intracellular GPCR signalling. We also present an alternative adapted bystander intracellular signalling assay for the GLP-1R in which we substitute the mini-G_s_ by a nanobody, Nb37, with specificity for active Gαs:GPCR complexes and no deleterious effect on the capacity for GLP-1R internalisation.

## Introduction

G protein-coupled receptors (GPCRs), the largest family of membrane proteins, orchestrate cellular responses to external stimuli, such as hormones and neurotransmitters (Rosenbaum et al., 2009). GPCRs signal through coupling to guanine nucleotide-binding regulatory proteins (G proteins) (Strader et al., 1994). Binding of an agonist induces a conformational change in the receptor that leads to activation of a trimeric G protein complex (Gahbauer and Bockmann, 2016; Strader et al., 1994), which occurs via by the exchange of guanosine diphosphate (GDP) by guanosine triphosphate (GTP) at the Gα subunit, plus dissociation of the Gβγ subunit (Daaka et al., 1998; Strader et al., 1994; Wolfgang et al., 1996). G protein activation subsequently enables a cascade of signal transduction events that leads to specific cellular effects (Strader et al., 1994). Once signal transmission has occurred, Gα subunits return to their inactive state through their intrinsic GTPase activity (Syrovatkina et al., 2016).

Considerable effort has been made in recent years to purify stable versions of GPCRs for structural analyses (Munk et al., 2019). One such approach consists in the co-expression of a GPCR with an engineered thermostable form of its cognate Gα subunit containing the GTPase domain, which has been termed a “mini-G protein” (Nehme et al., 2017). Mini-G proteins bind to and stabilise GPCRs in their active conformation and have therefore been widely used to generate structures from typically unstable active receptors (Nehme et al., 2017). In 2016, Carpenter & Tate developed a mini-G protein derived from Gαs, mini-G_s_ (Carpenter and Tate, 2016), which could stabilise the active β_1_-adrenergic receptor. Engineered mini-G proteins for other G protein families have since followed, including mini-G_olf_, mini-G_i1_, mini-G_o1_ and mini-G_q_ (Nehme et al., 2017; Wan et al., 2018). There are important differences between regular Gα proteins and their mini-G counterparts, including a) a shorter N-terminus which cannot interact with the Gβγ subunit; b) the insertion of a linker substituting the N-terminal α-helical domain; c) the removal of ten amino acid residues, leading to better *in vitro* stability; and d) the inclusion of a C-terminus mutation of the α5 helix to stabilise the GPCR:mini-G protein complexes (Nehme et al., 2017; Wan et al., 2018).

Unlike full-length Gα subunits, mini-G proteins are thought to be predominantly cytosolic under basal conditions, and are recruited to active receptors at different subcellular locations when stimulated by an appropriate agonist. Due to this ability of mini-G proteins to act as “conformational biosensors” for active GPCRs, many researchers have constructed vectors encoding a range of mini-G proteins fused to fluorescent or luminescent tags for in-cell experiments (Carpenter, 2018; Martemyanov and Garcia-Marcos, 2018; Wan et al., 2018). Examples include bioluminescence resonance energy transfer (BRET) and NanoLuc Binary Technology (NanoBiT) complementation assays for quantification of subtype-specific coupling to individual receptors (Wan et al., 2018) and spatiotemporal measurements of subcellularly-localised GPCR signalling (Crilly et al., 2021; Truong et al., 2021).

As the use of mini-G-based experimental approaches continues to expand, there is a need to carefully define the potential effects, if any, of mini-G co-expression on GPCR trafficking and signalling signatures from intracellular compartments to avoid hypothetical misinterpretation of results. With this in mind, we have examined here the effects of different mini-G subtypes on the β-arrestin-2 recruitment and trafficking profiles of the glucagon-like peptide-1 receptor (GLP-1R), a Gαs- and Gαq-coupled class B GPCR which plays important roles in the regulation of blood glucose levels and appetite (Baggio and Drucker, 2007; Sloop et al., 2018), as well as other prototypical GPCRs spanning the Gα subtype selectivity spectrum including the gastric inhibitory polypeptide receptor (GIPR) (Baggio and Drucker, 2007; Ismail et al., 2016), β_2_-adrenergic receptor (β2AR) (Bang and Choi, 2015), μ-opioid-receptor (μ-OR) (Pasternak and Pan, 2013), and free fatty acid receptor 2 (FFA2R) (Nilsson et al., 2003).

Our results unveil a previously unidentified effect of co-expressing mini-G proteins in specifically halting internalisation of their cognate GPCRs following agonist stimulation. This effect, which is most pronounced for Gαs-coupled receptors, is accompanied in the case of the GLP-1R by a complete inhibition of β-arrestin-2 recruitment despite near-normal levels of GRK2 recruitment, receptor clustering and segregation to specific lipid nanodomains. This poses important implications for the use of mini-G derivatives for the study of GPCR subcellular signal compartmentalisation, and should be taken into account in any experimental setup involving co-expression of modified cognate mini-G proteins to study GPCR behaviours. Alternative approaches to study signal compartmentalisation may include the use of nanobodies specific to active G protein-GPCR pairs, as demonstrated here for the GLP-1R with a bystander NanoBiT complementation assay based on a modified version of active Gαs-specific Nb37.

## Materials and Methods

### Cell culture

INS-1 832/3 cells (a gift from Professor Christopher Newgard, Duke University, USA) were cultured in RPMI 1640 medium (Gibco) supplemented with 10% FBS, 1% penicillin/streptomycin, 10 mM HEPES buffer, 100 mM sodium pyruvate, and 0.0004% β-mercaptoethanol. The derivative INS-1 832/3 cell lines used in this study were: INS-1 832/3 GLP-1R ^-/-^, INS-1 832/3 GIPR ^-/-^ [both kind gifts from Jackeline Naylor, Astra Zeneca (Naylor et al., 2016)], and INS-1 832/3 stably expressing N-terminally SNAP-tagged human GLP-1R and human GIPR (made in house from INS-1 832/3 GLP-1R^-/-^ or GIPR^-/-^ cells with G418 selection and FACS for SNAP-positive subpopulations). HEK293 cells were cultured in DMEM (Gibco) supplemented with 10% FBS and 1% penicillin/streptomycin. The HEK293 derivative cell lines used here were: HEK293 T-REx SNAP_f_-GLP-1R-SmBiT (Fang et al., 2020), HEK293 T-REx SNAP-FFA2R (generated in house by co-transfection of SNAP-tagged human FFA2R and pOG44 into HEK293 T-REx cells followed by hygromycin selection), and HEK293 SNAP-β_2_AR, SNAP-GIPR or SNAP-μ-OR (generated in house using the appropriate N-terminally SNAP-tagged GPCR with G418 selection as above). All cell lines were kept in a humidified incubator at a temperature of 37°C and a 5% CO_2_ atmosphere, and expression of SNAP-tagged receptors in T-REx lines was induced by addition of 0.1 μg/mL tetracycline 24 hours prior to performing experiments.

### Agonists

Peptide agonists were supplied by Wuxi Apptec at >90% purity. Isoproterenol, sodium proprionate, EGF and DAMGO were obtained from Sigma Aldrich. GLP-1-FITC has been described previously (Jones et al., 2017).

### Transfections

Transient transfections of Venus-, LgBiT- or NLuc-tagged mini-G constructs (all gifts from Professor Nevin Lambert, Augusta University, USA), Nb37-GFP (a gift from Professor Roshanak Irannejad, University of California San Francisco, USA), Gαs-YFP (Addgene plasmid #55781, a gift from Dr Catherine Berlot) and plasmids for the NanoBiT complementation and NanoBRET assays (see below) were performed using Lipofectamine 2000 (Thermo Fisher) for INS-1 832/3 and HEK293 cells according to the manufacturer’s instructions.

### Receptor internalisation assays

#### High-content microscopy internalisation assays

HEK293 cell lines stably expressing SNAP-tagged receptors of interest were transfected in 12-well plates with 0.1 μg mini-G_s_-, mini-G_i_- or mini-G_q_-Venus, or an mVenus plasmid (Addgene plasmid #27794, a gift from Professor Steven Vogel) as a negative control, plus 0.9 μg pcDNA 3.1 as carrier DNA. Cells were transferred to black, clear-bottom 96-well imaging plates (Falcon) coated with 0.01% poly-D-lysine 18 hours before the assay. Cells were treated for 15 minutes at 37°C with 1 μM BG-SS-649 (Marzook et al., 2021) (a gift from Dr Ivan Corrêa Jr, New England Biolabs) in complete media to reversibly label surface SNAP-tagged receptor populations. After washing 3 times, cells were treated in serum-free media ± agonist for 30 minutes at 37°C to induce receptor internalisation. Agonist was then removed, and cells were exposed to alkaline TNE buffer (pH 8.6) containing (or not) 100 mM Mesna for 5 minutes at 4°C to cleave BG-SS-649 bound to residual surface receptor. Microplates were then imaged using a Nikon Ti2E widefield microscope with LED light-source and 20X / 0.75 NA objective, with several transmitted phase contrast, Venus and BG-SS-649 epifluorescence images acquired per well using appropriate filters. Image analysis was performed using Fiji: cell-containing regions were first segmented using PHANTAST (Jaccard et al., 2014), after which Venus-expressing and non-expressing cells were separately segmented. After flat-field correction using BaSiC (Peng et al., 2017), background-subtracted BG-SS-649 intensities in the above cell populations were quantified, and agonist-mediated internalisation determined by subtracting the average intensity from non-agonist-treated but Mesna-exposed cells, indicative of constitutive receptor internalization and non-specific probe uptake. In some cases, a further normalisation was applied by expressing the agonist-induced internalisation signal relative to total BG-SS-649 labelling measured in cells not exposed to Mesna (Marzook et al., 2021). This is equivalent to “percentage internalisation” but, as we observed that internalised BG-SS-649 typically displays higher signal intensity than when at the surface, >100% internalisation can sometimes be recorded using this method; we therefore refer to this quantification as “apparent percentage internalisation”.

#### Time-lapse microscopy internalisation assays

INS-1 832/3 cells stably expressing SNAP-GLP-1R or SNAP-GIPR and HEK293 or HEK293 T-REx cells stably expressing SNAP-β2AR, −μ-OR or - FFA2R were seeded and transfected in 24-well plates with 0.05 μg mini-G_s_-, mini-G_i_- or mini-G_q_-Venus, or mVenus as a negative control. SNAP_f_-EGFR (New England Biolabs) was transiently co-transfected into INS-1 832/3 cells with each of the above plasmids. Additionally, INS-1 832/3 SNAP-GLP-1R cells were transfected with 0.05 μg Nb37-GFP or empty GFP in a parallel experiment. Separately, INS-1 832/3 SNAP-GLP-1R cells were also transfected with 0.05 μg Gαs-YFP. 24 hours post-transfection, the cells were transferred to MatTek dishes (MatTek Life Sciences) and left to adhere overnight. The next day, the cells were labelled with 1 μM SNAP-Surface 549 (New England Biolabs) for 15 minutes at 37°C. For SNAP_f_-EGFR, cells were preincubated for 1 hour in serum-free medium supplemented with 0.1% BSA prior to labelling. Following labelling, the cells were washed once with 1X PBS and imaged in RPMI without phenol red (ThermoFisher). Fluorescence was recorded for 1 minute, after which the cells were stimulated with the agonist for its cognate receptor and time-lapse recordings taken each 6 seconds for a further 10 minutes at 37°C in a Nikon Eclipse Ti spinning disk microscope with temperature control coupled to a 16-bit, 512×512 pixel back illuminated EM-CCD camera [ImageEM 9100-13 (Hamamatsu)] using a 60X oil immersion objective. GLP-1, GIP, and isoproterenol were used at 100 nM, DAMGO was used at 1 μM, proprionate at 1 mM and EGF at 100 μg/mL. Data was analysed in Fiji: three separate lines were drawn across the membranes of each cell and the intensity calculated using a macro designed in house by the Facility for Imaging by Light Microscopy (FILM) at Imperial College London, UK. Intensities were normalised to average baseline reading for each video and used to calculate the percentage of receptor at the plasma membrane. These values were then transformed to display the percentage of receptor internalised over a period of 10 minutes.

### Mini-G plasma membrane recruitment NanoBRET assay

HEK293 cell lines stably expressing SNAP-tagged receptors were transfected in 12-well plates with 0.1 μg NLuc-tagged mini-G_s_, mini-G_q_ or mini-G_i_, plus 0.1 μg KRAS-Venus (a gift from Professor Nevin Lambert, Augusta University, USA) and 0.8 μg pcDNA 3.1. After 24 hours, cells were detached, seeded into white 96-well half area plates in HBSS and stimulated with agonist or vehicle for 30 minutes at 37°C. Furimazine (1:20 dilution, Promega) was then added and the plate promptly transferred to a Flexstation 3 plate reader at 37°C. Luminescence was recorded over 10 minutes at 460 nm and 535 nm in parallel, with the average signal at each wavelength used to calculate a 535/460 BRET ratio. Agonist-induced BRET was calculated by subtracting the average signal from vehicle wells, and further normalised to cell surface expression levels obtained in the same cell lines by high-content microscopy as above.

### Additional NanoBiT and NanoBRET assays

Specific assay components are described below, but for all assays, 24 hours after transfection cells were detached, resuspended in NanoGlo Live Cell Reagent (Promega) with furimazine (1:20 dilution) and seeded into white 96-well half area plates. Luminescence was recorded at 37°C in a Flexstation 3 plate reader, with total luminescent signal used for NanoBiT assays and dual wavelength acquisition (460 and 535 nm) for NanoBRET assays. A 5-minute baseline recording was followed by agonist addition and serial measurements over 30 minutes; readings were taken every 30 seconds and normalised to well baseline and then average vehicle-induced signal was subtracted to establish the agonist-induced effect. Areas under the curve (AUC) were calculated for each concentration of agonist and fitted to four-parameter curves using Prism (GraphPad Software).

#### β-arrestin-2 recruitment NanoBiT assays

HEK293 T-REx SNAP_f_-GLP-1R-SmBiT cells were seeded in 12-well plates and co-transfected with 0.1 μg β-arrestin-2 fused at the N-terminus to LgBiT (LgBiT-β-arrestin-2; Promega, plasmid no. CS1603B118) and either 0.1 μg mini-G_s_-Venus or mVenus supplemented with 0.8 μg pcDNA 3.1.

#### Nb37-based bystander NanoBiT assays

Nb37 cDNA (synthesised by GenScript with codon optimisation) was C-terminally fused to SmBiT with a 15-amino-acid flexible linker (GGSGGGGSGGSSSGGG), and the resulting construct referred to as Nb37-SmBiT. The C-terminal KRAS CAAX motif (SSSGGGKKKKKKSKTKCVIM) was N-terminally fused with LgBiT (LgBiT-CAAX). The Endofin FYVE domain (amino-acid region Gln739-Lys806) was C-terminally fused with LgBiT (Endofin-LgBiT). Gαs (human, short isoform), Gβ1 (human), Gγ2 (human), and RIC8B (human, isoform2) plasmids were produced in house. These constructs were inserted into pcDNA 3.1 or pCAGGS expression plasmid vectors. HEK293 cells were seeded in 6-well plates and co-transfected with 0.2 μg SNAP-GLP-1R, 0.5 μg Gαs, Gβ1, and Gγ2, 0.1 μg RIC8B, 0.1 μg CAAX-LgBiT or 0.5 μg Endofin-LgBiT with 0.1 μg or 0.5 μg Nb37-SmBiT, respectively.

#### Subcellular receptor localisation NanoBRET assays

HEK293 cells were seeded in 12-well plates and co-transfected with 0.5 μg each of SNAP-GLP-1R-NLuc (generated in house) and KRAS- or Rab5-Venus (a gift from Professor Nevin Lambert, Augusta University, USA).

#### GRK2 recruitment NanoBRET assay

HEK293 cells were seeded in 12-well plates and co-transfected with 0.05 μg GRK2-Venus (a gift from Professor Denise Wootten, Monash University, Australia) and 0.05 μg SNAP-GLP-1R-NLuc, plus either 0.1 μg mini-G_s_-LgBiT supplemented with 0.8 μg pcDNA 3.1 or 0.9 μg pcDNA 3.1.

### Membrane fractionation for lipid raft segregation assays

INS-1 832/3 SNAP GLP-1R cells were seeded into 60-mm dishes and transfected with either 3.4 μg of mini-G_s_-Venus or mVenus as a control. Cells were treated with either vehicle or 100 nM GLP-1 for 2 minutes at 37°C. Following stimulation, cells were washed with PBS and osmotically lysed in 20 mM Tris-HCl, pH 7, with 1% protease inhibitor (Roche) and 1% phosphatase inhibitor cocktail (Sigma-Aldrich). Cells were then passed through a 21-gauge needle and ultracentrifuged at 41,000 rpm at 4°C for 1 hour. Following this, pellets were resuspended in PBS supplemented with 1% Triton-X100 plus 1% protease inhibitor and 1% phosphatase inhibitor cocktail and rotated at 4°C for 30 minutes. Samples were then ultracentrifuged again at 41,000 rpm at 4°C for 1 hour. The detergent soluble membrane fractions (supernatant, DSMs) were separated from the detergent resistant membrane fractions (pellet, DRMs), which were then resuspended in 1% SDS supplemented with protease and phosphatase inhibitors. The membrane fractions were then sonicated, centrifuged at 13,000 rpm at 4°C for 5 minutes and supernatants collected. Samples were resolved by SDS-PAGE in urea loading buffer and analysed by Western blot against the SNAP-tag (rabbit polyclonal anti-SNAP-tag antibody P9310S, New England Biolabs) to detect levels of SNAP-GLP-1R in the different fractions, as well as flotillin (anti-flotillin-1 antibody, 849802, BioLegend) as a DRM marker (Salzer and Prohaska, 2001), to assess the quality of the fractionation and as a loading control. Blots were developed with the Clarity Western enhanced chemiluminescence (ECL) substrate system (BioRad) in a Xograph Compact X5 processor and specific band densities quantified in Fiji. SNAP-GLP-1R content in the DRM fractions for the different conditions was normalised to that of flotillin.

### Receptor co-localisation assays

SNAP-GLP-1R subcellular localisation, Nb37-GFP co-localisation into endosomes and time-lapse microscopy for SNAP-GLP-1R and Rab5-mCherry co-localisation (Addgene plasmid # 49201, a gift from Dr Gia Voeltz) were performed by confocal microscopy in cells previously labelled with the indicated SNAP-Surface fluorescent probe (New England Biolabs) using a Zeiss LSM-780 microscope with a 63X / 1.4 NA oil objective from the FILM Facility, Imperial College London, UK.

### Lipid nanodomain cAMP FRET assays

A-kinase-anchoring proteins (AKAPs) are scaffold proteins in charge of recruiting PKA at specific cAMP activation sites (Dodge and Scott, 2000). AKAP79 is one such scaffold compartmentalised at lipid nanodomains by virtue of its palmitoylation (Delint-Ramirez et al., 2011; Dodge and Scott, 2000; Woolfrey et al., 2015). Fluorescence resonance energy transfer (FRET) using a cAMP biosensor based on AKAP79, AKAP79-CUTie (Surdo et al., 2017) (a gift from Professor Manuela Zaccolo, University of Oxford, UK) was carried out to quantify cAMP generation specifically from these nanodomains. HEK293 T-REx SNAP_f_-GLP-1R-SmBiT cells were transfected with 0.5 μg AKAP79-CUTie and either 0.5 μg of mini-G_s_-LgBiT or 0.5 μg pcDNA 3.1. The next day, cells were seeded onto 96-well imaging plates in HBSS and a baseline FRET reading was taken before stimulation with a range of GLP-1 concentrations. FRET values were read on a Flexstation 3 plate reader for 30 minutes with the following settings: λex = 440 nm, λem = 485 and 535 nm. FRET was quantified as the ratio of fluorescent signal at 535 nm to that at 485 nm after subtraction of background signal at each wavelength to generate concentration responses which were fitted to four-parameter curves using Prism as above. Samples were lysed at the end of the assay and analysed for total cAMP using HTRF (Cisbio cAMP Dynamic assay).

### Clustering assays

The assays were performed as previously described (Buenaventura et al., 2019). INS-1 832/3 SNAP-GLP-1R cells were transfected with either 1 μg mini-G_s_-LgBiT or pcDNA 3.1. 24 hours post-transfection the cells were dual-labelled with the time-resolved (TR)-FRET probe SNAP-Lumi4-Tb (40 nM, Cisbio) and SNAP-Surface 647 (1 μM, New England Biolabs) for 60 minutes at room temperature. Cells were washed and placed in HBSS in a white 96-well plate for a 10-minute baseline measurement at 37 °C using a Spectramax i3x (Molecular Devices) plate reader fitted with a TR-FRET filter set (λEx = 335 nm, λEm = 616 nm and 665 nm, delay 30 μs, integration time 400 μs). TR-FRET was sequentially measured after agonist addition. The ratio of fluorescent signals at both emission wavelengths (665/616) was considered indicative of clustering as it reflects both transient and stable interactions between receptor protomers occurring within the long fluorescence lifetime of the excited terbium cryptate.

### Plasma membrane AP2 hotspot co-localisation assay

HEK293 T-REx SNAP_f_-GLP-1R cells were co-transfected with an AP2-HA plasmid (μ2-HA-WT; Addgene plasmid # 32752, a gift from Professor Alexander Sorkin) and either mini-G_s_-Venus or mVenus as a control. 24 hours post-transfection, cells were transferred to MatTek dishes and left to adhere overnight. The following day, cells were stimulated with GLP-1 for 2 minutes, fixed and processed by immunofluorescence with an anti-HA monoclonal antibody (clone HA-7, Sigma Aldrich) and a secondary anti-mouse AlexaFluor 555 (Thermo Fisher) and left unmounted in PBS. Co-localisation with plasma membrane AP2 hotspots was performed by total internal reflection fluorescence (TIRF) microscopy with the Nikon Eclipse Ti spinning disk microscope coupled to a TIRF iLas2 module to control laser angle (Roper Scientific), and a Quad Band TIRF filter cube (TRF89902, Chroma) using a 100X / 1.49 NA oil immersion TIRF objective.

### Statistical Analyses

All statistical analyses were performed using GraphPad Prism 9.1.2 (GraphPad Software). All values are given as mean ± SEM and statistical significance was calculated using either Student’s t-test (two-tailed) when comparing two variables or one-way ANOVA for comparison between more than two variables, unless stated otherwise.

## Results

This project stems from an initial observation of strikingly sustained mini-G_s_ retention at the plasma membrane after GLP-1R stimulation, which appears incompatible with the previously known fast internalisation kinetics of this receptor (Shaaban et al., 2016). Specifically, in GLP-1R-expressing HEK293 cells, we have previously observed that NanoLuc-tagged mini-G_s_ remains associated with the plasma membrane after an initial agonist-mediated recruitment phase, contrasting with the rapid disappearance of NanoLuc-tagged GLP-1R from the cell surface measured in parallel in the same cell system, but without mini-G_s_ co-expression (Lucey et al., 2021).

With this in mind, we performed recruitment experiments with mini-G_s_-NLuc co-expressed with either the plasma membrane KRAS-Venus or the endosomal Rab5-Venus reporters in HEK293 cells stably expressing SNAP-GLP-1R (Figure 1A and B). Similarly to our previous observations (Lucey et al., 2021), whilst we could clearly detect mini-G_s_ plasma membrane recruitment following GLP-1 stimulation, the pattern of recruitment was sustained and did not subside over time, as would be expected from the known kinetics as well as our own examination of GLP-1R internalisation times (Figure 1C). This effect correlated with an almost negligible detection of mini-G_s_ recruitment to the early endosomal compartment (Figure 1A and B). When this experiment was performed in SNAP-GLP-1R-expressing INS-1 832/3 beta cells, a cell type which normally expresses the GLP-1R endogenously, we could again detect prolonged mini-G_s_ recruitment to the plasma membrane (albeit with a slow tendency to decrease) (Figure 1D), despite a much more rapid and almost complete GLP-1R internalisation to Rab5-positive endosomes following receptor activation when examined without mini-G_S_ expression (Figure 1E, Supplementary Movie 1). Moreover, we were unable to detect any mini-G_s_ recruitment to endosomes in these cells (not shown). These results could reasonably be interpreted as demonstrating low or non-existent levels of active GLP-1Rs in endosomes, particularly in beta cells, which is surprising given the existence of a number of previous reports of endosomal GLP-1R cAMP signalling using alternative approaches (Girada et al., 2017; Kuna et al., 2013).

**Figure 1:**
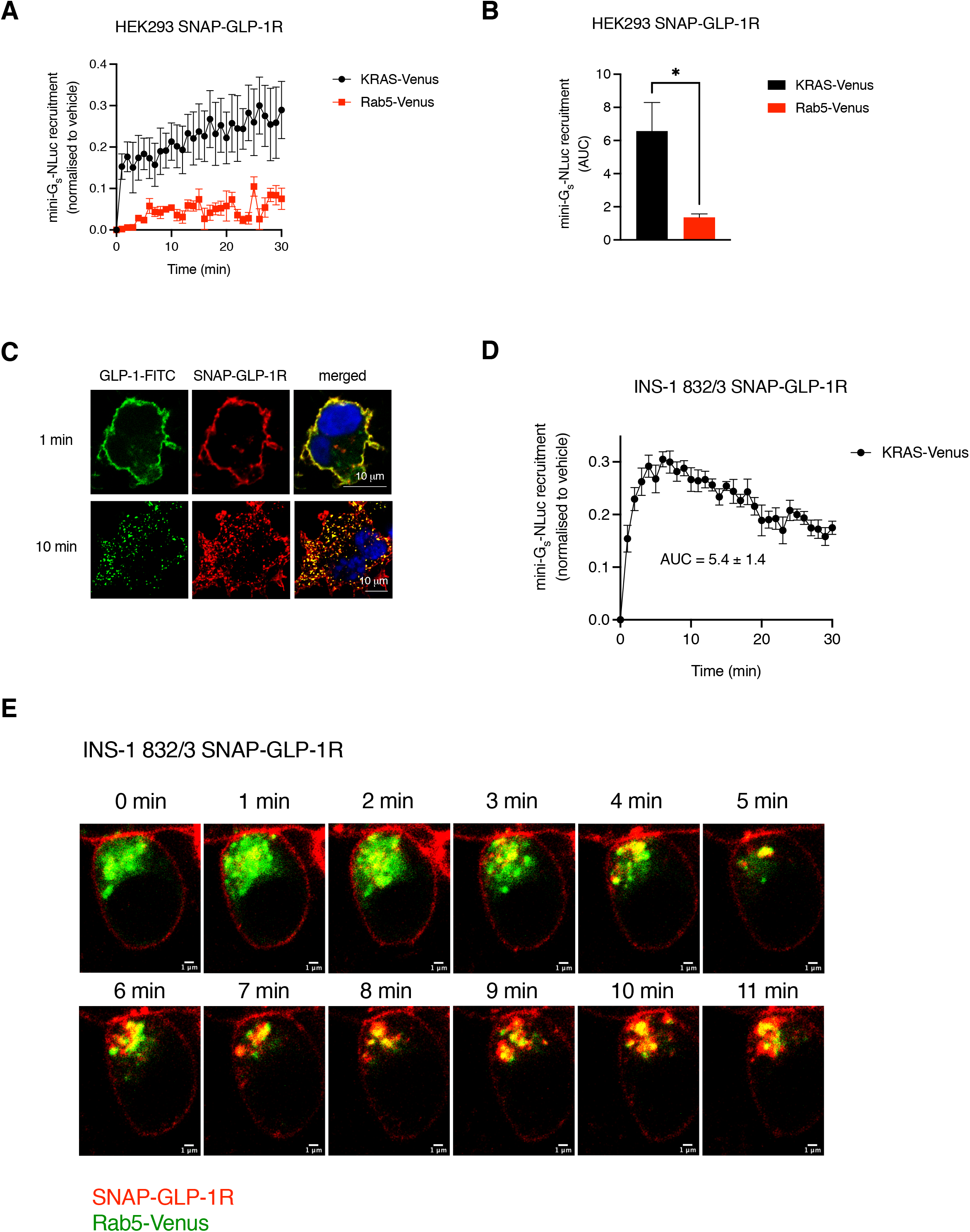
Disconnect between mini-Gs plasma membrane recruitment and GLP-1R internalisation. (**A**) Bystander NanoBRET assay measuring miniG_s_-NLuc recruitment to the plasma membrane (with KRAS-Venus) *vs* endosomes (with Rab5-Venus) for 30 min in response to 100 nM GLP-1 in HEK293 SNAP-GLP-1R cells; *n*=5. (**B**) AUC quantification from (A). (**C**) Subcellular localisation of SNAP-GLP-1R (labelled with SNAP-Surface 549, red) after 1 min and 10 min of 100 nM GLP-1-FITC (green) stimulation in HEK293 SNAP-GLP-1R cells by confocal microscopy; nucleus (DAPI), blue; size bars as indicated. (**D**) Bystander NanoBRET assay of miniG_s_-NLuc recruitment to the plasma membrane as in (A) but in INS-1 832/3 SNAP-GLP-1R cells; *n*=4. AUC as indicated. (**E**) Time-lapse confocal microscopy analysis of SNAP-GLP-1R (labelled with SNAP-Surface 549, red) trafficking to Rab5-Venus (green)-positive endosomes in INS-1 832/3 SNAP-GLP-1R cells. Data is mean ± SEM; *p<0.05 by paired t-test.

These puzzling results prompted us to analyse the potential effects of co-expressing a range of mini-G subtypes on SNAP-GLP-1R internalisation rates. Using mini-G fused to a fluorescent Venus marker to allow analysis of responses in mini-G-expressing *versus* non-expressing cells in the same field of view, we found that agonist-induced SNAP-GLP-1R internalisation was severely blunted by mini-G_s_ expression in HEK293 cells, with a smaller inhibitory effect of mini-G_q_, and no effect of mini-G_i_ or mVenus alone (Figure 2A and B). This rank order correlated with the degree of GLP-1-mediated recruitment of each mini-G subtype to the plasma membrane, as measured by NanoBRET (Figure 2C). Time-lapse microscopy experiments in stable SNAP-GLP-1R-expressing INS-1 832/3 pancreatic beta cells showed exactly the same pattern (Figure 2D–F), indicating that this effect of mini-G expression is also observed in the native cellular environment of the GLP-1R. However, inhibition of GLP-1R internalisation was not observed when a YFP-tagged full-length Gαs protein was co-expressed, suggesting that this effect is restricted to its mini-G_s_ counterpart (Supplementary Figure 1A and B). Additionally, co-expression of mini-G_s_ also did not impact on the GLP-1R capacity for cAMP generation (Supplementary Figure 1C). These results offer a potential explanation for our previous failure to detect endosomal GLP-1R signalling, as a mini-G_s_-mediated halt of receptor trafficking would impede proper quantification of signalling from this intracellular location.

**Figure 2:**
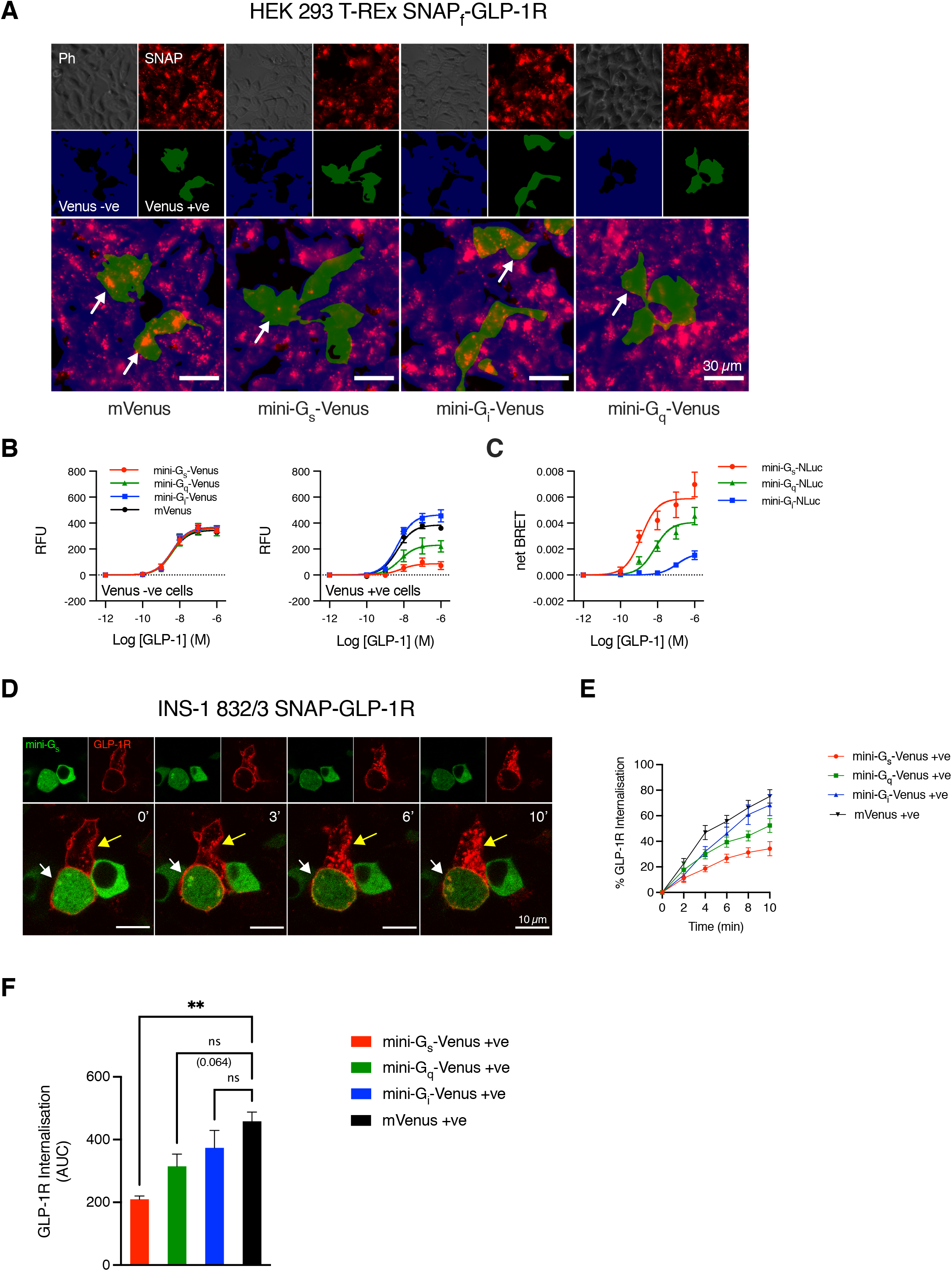
Direct correlation between subtype mini-G-induced inhibition of internalisation and degree of coupling to the GLP-1R. (**A**) Representative images showing internalised SNAP_f_-GLP-1R after treatment with 100 nM GLP-1 for 30 min in HEK293 T-REx SNAP_f_-GLP-1R-SmBiT cells co-transfected with the indicated Venus construct. Residual surface label was then removed using Mesna. Phase contrast (Ph) and BG-SS-649 (SNAP; red) epifluorescence images are shown, along with segmented regions indicating Venus-expressing (green) and non-expressing (blue) cells. The white arrows highlight how mini-G_s_ and mini-G_q_-expressing cells typically show reduced levels of internalised GLP-1R. (**B**) Quantification of experiment shown in (A), indicating average fluorescence of internalised GLP-1R in Venus-expressing and non-expressing cells treated with a range of concentrations of GLP-1; *n*=5 independent acquisitions. (**C**) Recruitment of each NLuc-tagged mini-G subtype to the plasma membrane in HEK293 T-REx SNAP_f_-GLP-1R-SmBiT cells co-transfected with KRAS-Venus and stimulated with GLP-1 for 30 min; *n*=5. (**D**) Time-lapse spinning disk images of SNAP-GLP-1R (labelled with SNAP-Surface 549, red) subcellular localisation from a representative mini-G_s_-Venus positive (green; white arrow) *vs* negative (yellow arrow) INS-1 832/3 SNAP-GLP-1R cell in response to 100 nM GLP-1 at the indicated frame times; size bars as indicated. (**E**) Percentage of GLP-1R internalisation in Venus-positive INS-1 832/3 SNAP-GLP-1R cells expressing each of the different mini-G subtypes tested (mini-G_s_, -G_q_ and G_i_) as well as empty mVenus as a control; *n*=3 independent time-lapse acquisitions. (**F**) AUC quantifications from (E). Data is mean ± SEM; **p<0.01 by one-way ANOVA with Dunnett’s test; ns (non-significant).

Following G protein activation, GPCRs are Ser/Thr phosphorylated by G protein receptor kinases (GRKs). The phosphorylated receptor is then prompted to interact with β-arrestins, resulting in G protein uncoupling, receptor recruitment to clathrin-coated pits (CCPs) and internalisation (Hanyaloglu, 2018). Given its apparent effects on GLP-1R internalisation, we next asked whether mini-G_s_ expression would also alter recruitment of its preferred GRK and β-arrestin isoforms, namely GRK2 and β-arrestin-2 (Figure 3). Co-expression of mini-G_s_ *versus* an empty pcDNA 3.1 construct led to an almost complete inhibition of agonist-mediated β-arrestin-2 recruitment to the GLP-1R (Figure 3A and B), whilst GRK2 recruitment showed a non-significant trend towards reduction (Figure 3C and D). This suggests that the stability of the mini-G_s_:GLP-1R interaction is sufficient to prevent its displacement by β-arrestins, despite near normal GRK2 recruitment.

**Figure 3:**
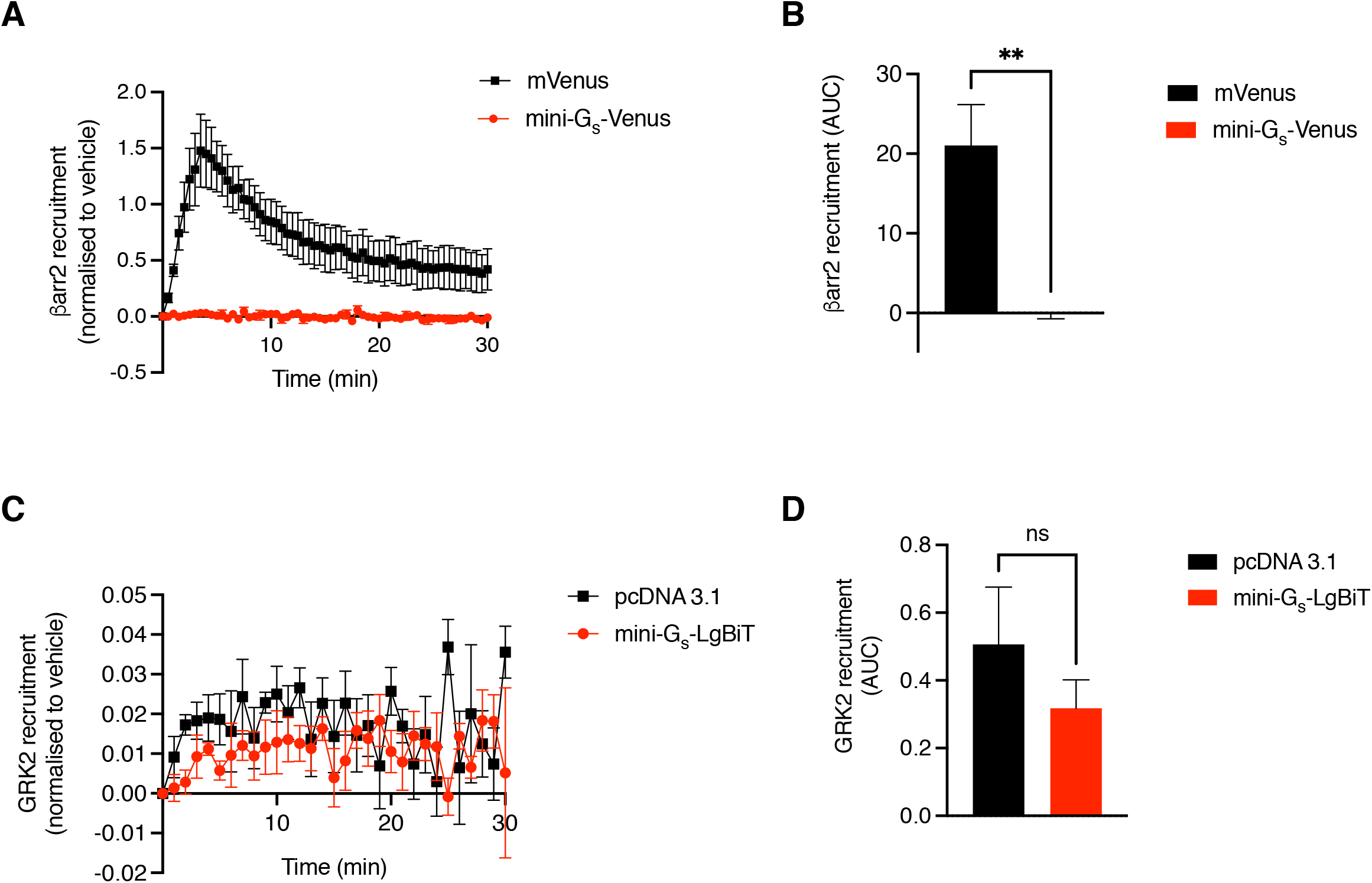
Effect of mini-Gs co-expression on GLP-1R capacity to recruit β-arrestin-2 and GRK2 after GLP-1 stimulation. (**A**) NanoBiT assay measuring β-arrestin-2-LgBiT recruitment to SNAP_f_-GLP-1R-SmBiT for 30 min in response to 100 nM GLP-1 in HEK293 T-REx SNAP_f_-GLP-1R-SmBiT cells co-expressing mini-G_s_-Venus or mVenus as a control; *n*=5. (**B**) AUC quantification from (A). (**C**) NanoBRET assay measuring GRK2-Venus recruitment to SNAP-GLP-1R-NLuc for 30 min in response to 100 nM GLP-1 in HEK293 cells co-expressing mini-G_s_-LgBiT or pcDNA 3.1 as a control; *n*=5. (**D**) AUC quantification from (C). Data is mean ± SEM; **p<0.01 by paired t-test; ns (non-significant).

We have previously shown that the biologically active form of the GLP-1R is preferentially clustered into and signals from cholesterol-rich plasma membrane nanodomains prior to its internalisation into the endocytic pathway (Buenaventura et al., 2019). We therefore next checked whether mini-G_s_ co-expression could interfere with this membrane compartmentalisation process (Figure 4). In INS-1 832/3 SNAP-GLP-1R-expressing cells, we found that mini-G_s_ co-expression had no effect either on the capacity for clustering of the GLP-1R (Figure 4A and B) or its recruitment to lipid nanodomains (Figure 4C–E) following GLP-1 stimulation, suggesting that active mini-G_s_-bound GLP-1R retains its capacity to segregate to these plasma membrane hotspots. We did however find a small but significant increase in the amount of receptor localised to lipid rafts under vehicle conditions in the presence of mini-G_s_, suggesting the existence of some degree of mini-G_s_ binding to and stabilisation of the GLP-1R in an active conformation even in the absence of agonist. Consistent with these observations, we could not detect any significant differences in the degree of raft-specific cAMP generation, measured with the cAMP FRET biosensor AKAP79-CUTie in HEK293 T-REx SNAP_f_-GLP-1R cells in response to GLP-1 (Figure 4F, Supplementary Figure 2A). Additionally, normal recruitment of the receptor to lipid nanodomains and lipid raft cAMP generation was accompanied by co-localisation of mini-G_s_-Venus to AP2-rich plasma membrane hotspots, as assessed by TIRF microscopy of HEK293 T-REx SNAP_f_-GLP-1R cells following GLP-1 exposure (Supplementary Figure 2B), suggesting that the blockade in internalisation takes place very late in the CCP segregation and endocytosis process.

**Figure 4:**
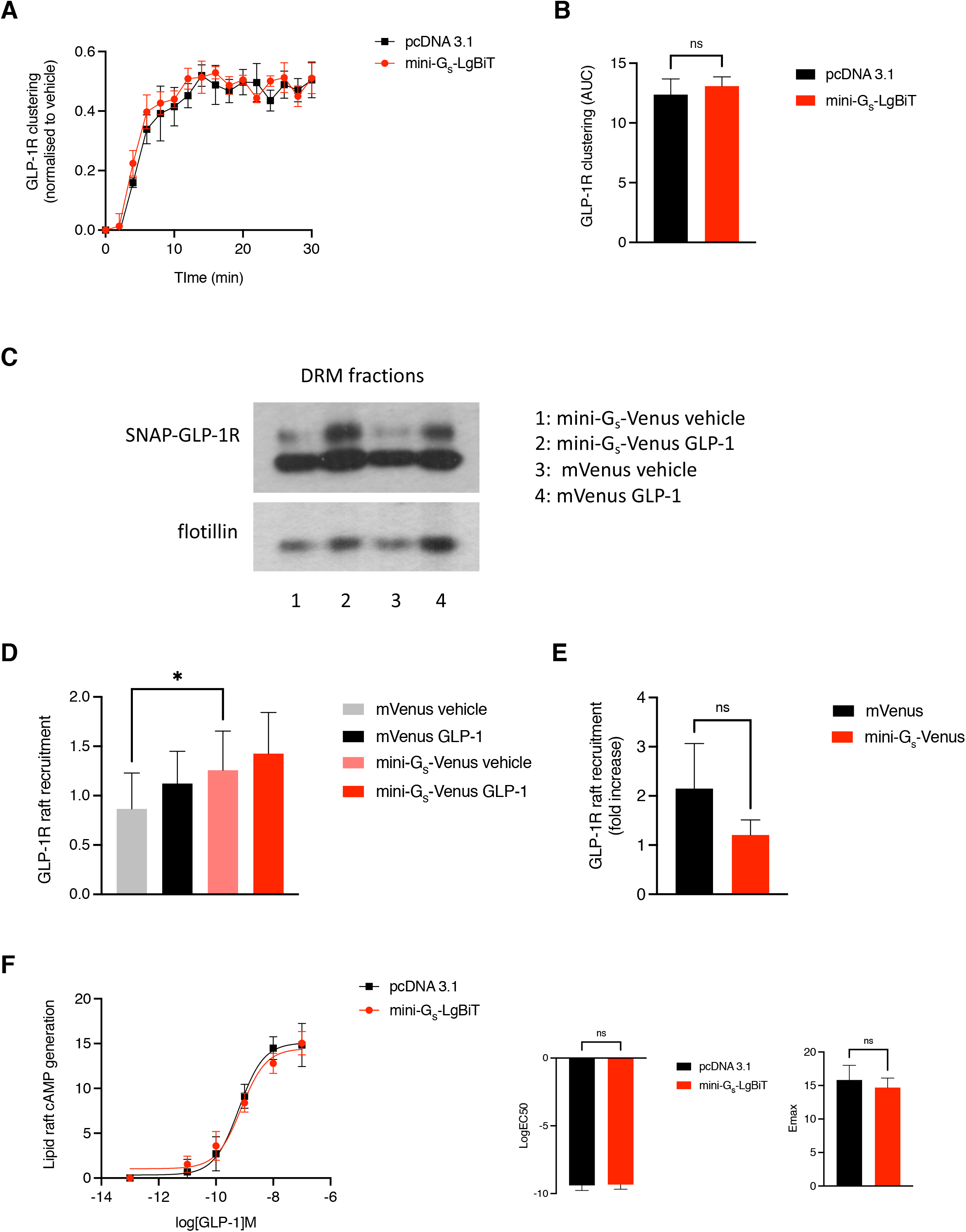
Effect of mini-Gs co-expression on the GLP-1R capacity for clustering and segregation to plasma membrane cholesterol-rich lipid nanodomains. (**A**) Clustering assay measured by TR-FRET in INS-1 832/3 SNAP-GLP-1R cells co-expressing mini-G_s_-LgBiT or pcDNA 3.1 as a control; *n*=5. (**B**) AUC quantification from (A). (**C**) Representative blots from detergent-resistant membrane (DRM) fractions collected from INS-1 832/3 SNAP-GLP-1R cells co-expressing mini-G_s_-Venus or mVenus as control under vehicle or 100 nM GLP-1-stimulated conditions. SNAP-GLP-1R detected with an anti-SNAP tag Ab and flotillin used as a DRM-specific loading control. (**D**) Quantification of SNAP-GLP-1R lipid raft recruitment from (C); *n*=5. (**E**) Lipid raft recruitment fold-increase to vehicle, calculated from (D). (**F**) Lipid raft-localised cAMP generation, dose responses to GLP-1 measured with AKAP79-CUTie in HEK293 T-REx SNAPf-GLP-1R-SmBiT cells co-expressing mini-Gs-LgBiT or pcDNA 3.1 as a control; logEC50 and Emax comparisons from the dose response curves displayed; *n*=6. Data is mean ± SEM; *p<0.05 and ns (non-significant) by paired t-test.

In view of the confounding effects of mini-G_s_ co-expression, we next devised an alternative bystander NanoBRET strategy in order to more reliably quantify both endosomal and plasma membrane origin signalling without inducing artefactual effects on receptor trafficking. For this we made use of a nanobody, Nb37, previously developed to bind to Gαs in complex with active GPCRs (Irannejad et al., 2013). Co-expression of a Nb37-GFP fusion protein had a minimal effect on the rate of internalisation of the GLP-1R (Figure 5A). Using Nb37 fused to SmBiT that we co-expressed with the plasma membrane and endosomal bystander reporters CAAX- and Endofin-LgBiT, respectively (Figure 5B and C) (McGlone et al., 2021), we could now detect a more expected pattern of Nb37 recruitment to the plasma membrane, whereby recruitment reached a peak after about 5-8 minutes of stimulation with GLP-1 and then steadily declined. In parallel, this method also allowed us to detect a robust increase in endosomal signalling by the receptor, validated by the presence of Nb37-GFP and SNAP-GLP-1R in Rab5-positive endosomes by confocal microscopy (Figure 5D).

**Figure 5:**
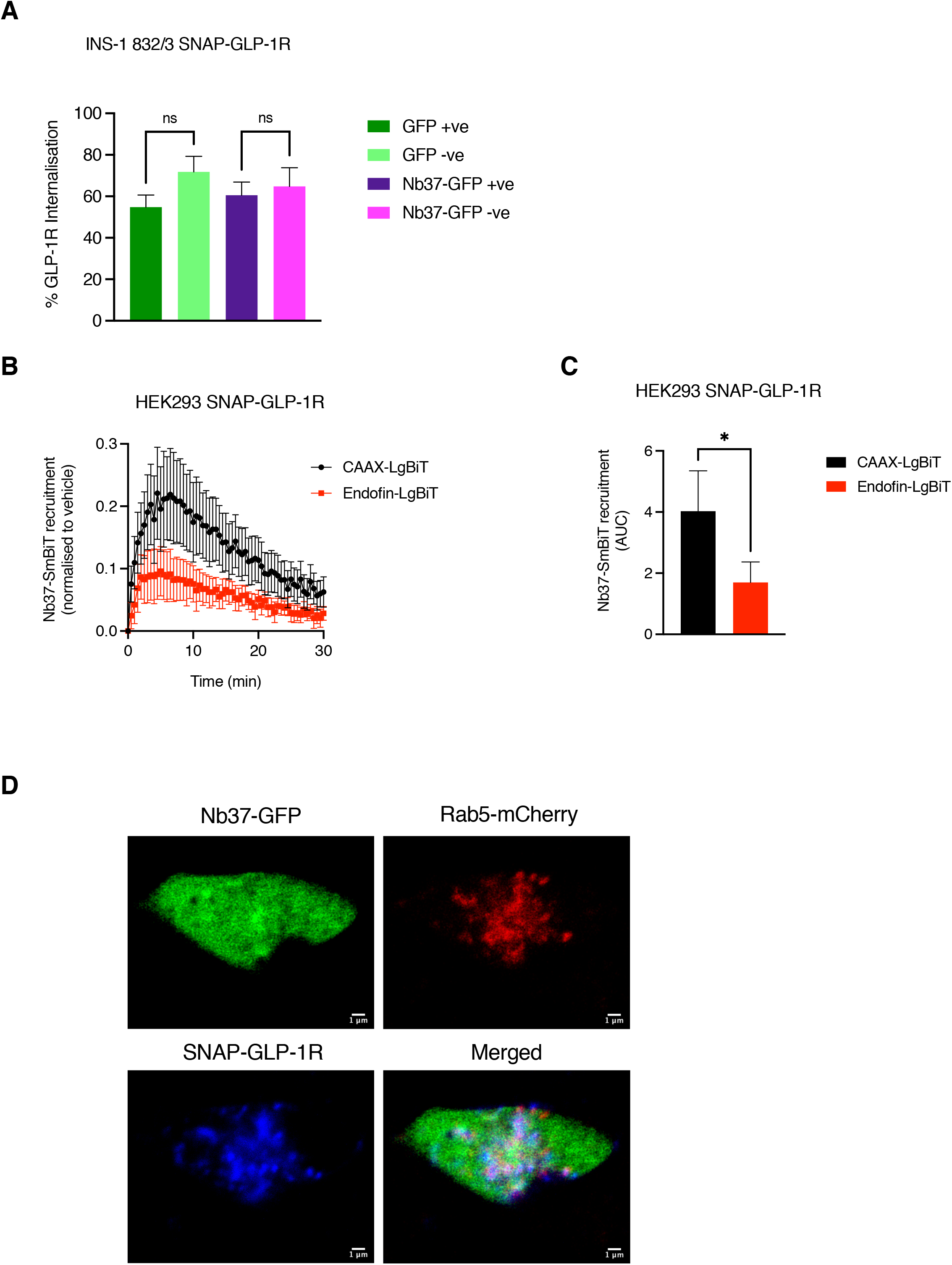
Effect of Nb37 co-expression on GLP-1R internalisation and Nb37-based bystander NanoBiT assay of GLP-1R subcellular signalling. (**A**) Percentage of GLP-1R internalisation in GFP-positive *vs* −negative INS-1 832/3 SNAP-GLP-1R cells expressing Nb37-GFP or an empty GFP vector as a control; *n*=4 independent experiments. (**B**) Bystander NanoBiT assay measuring Nb37-SmBiT recruitment to the plasma membrane (with CAAX-LgBiT) or to endosomes (with Endofin-LgBiT) for 30 min in response to 100 nM GLP-1 in HEK293 SNAP-GLP-1R cells; *n*=6. (**C**) AUC quantification from (B). (**D**) Subcellular localisation of Nb37-GFP (green) and SNAP-GLP-1R (labelled with SNAP-Surface 647, blue) to endosomes (labelled by co-expression of Rab5-mCherry, red) after 10 min stimulation with 100 nM GLP-1 in HEK293 SNAP-GLP-1R cells by confocal microscopy; size bars as indicated. Data is mean ± SEM; ns (non-significant) by one-way ANOVA with Dunnett’s test; *p<0.05 by paired t-test.

Finally, we asked whether the trafficking of other receptors beyond the GLP-1R could also be affected by mini-G protein co-expression. To this end, we determined the pattern of mini-G subtype recruitment by NanoBRET (Figure 6A), and the effect of mini-G protein expression on receptor internalisation by high content microscopy (Figure 6B), of the class B glucose-dependent insulinotropic polypeptide receptor (GIPR) and three class A GPCRs: the β2-adrenergic receptor (β2AR), the μ-opioid receptor (μ-OR) and the free fatty acid 2 receptor (FFA2R). As for the GLP-1R, we observed a clear correlation between the degree of inhibition of receptor internalisation and the selective coupling to each mini-G subtype, although the intensity of this effect was somewhat variable depending on each individual receptor (Figure 6C). The inhibition of internalisation results were corroborated by time-lapse confocal microscopy for each GPCR with the more prominently coupled mini-G subtype (Figure 6D–G). We also tested these effects on a non-GPCR, using the epidermal growth factor receptor (EGFR) as a model receptor tyrosine kinase (RTK), and concluded that expression of any mini-G subtype did not affect its degree of internalisation (Supplementary Figure 3).

**Figure 6:**
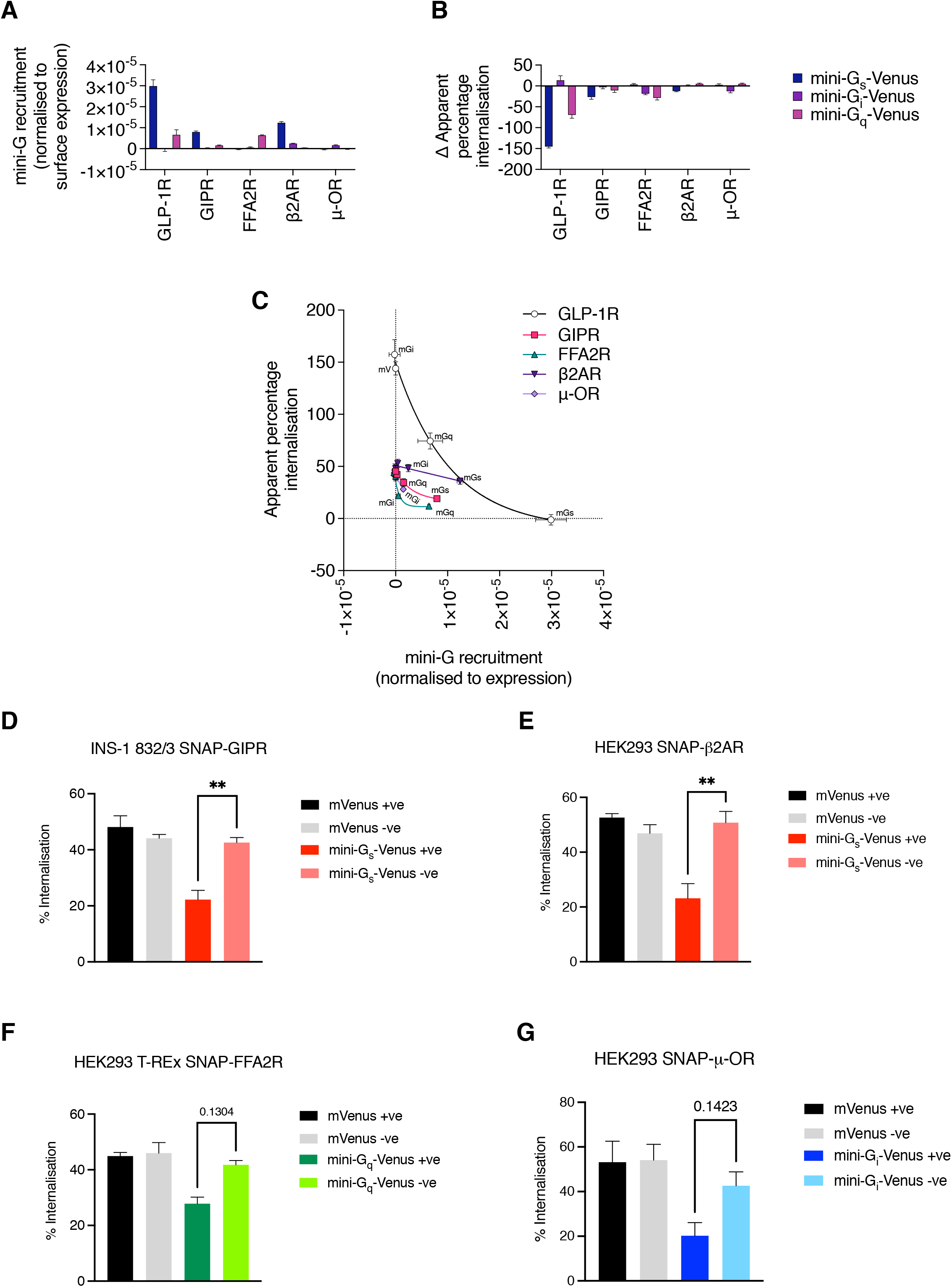
Effect of mini-G co-expression across different GPCR classes. (**A**) Recruitment of each NLuc-tagged mini-G subtype to the plasma membrane in HEK293 cell lines stably expressing the indicated GPCR, co-transfected with KRAS-Venus and stimulated with the appropriate cognate agonist (100 nM GLP-1, 100 nM GIP, 1 mM proprionate, 100 nM isoproterenol, or 1 μM DAMGO) for 30 min; *n*=5. The NanoBRET signal has been adjusted to account for differences in receptor expression as measured by high content microscopy. (**B**) Impact on receptor internalisation of co-expression with the indicated mini-G-Venus fusion construct, determined by high content microscopy after 30 min stimulation with the appropriate cognate agonist as in (A); *n*=5. The absolute difference between “apparent percentage internalisation” (see Methods) in mini-G *versus* mVenus control cells is shown. (**C**) The association between mini-G subtype recruitment (shown in A) and receptor internalisation in Venus-expressing cells (same experiment as in B) for each GPCR. (**D**) Quantification of GIPR internalisation in Venus-positive *vs* −negative INS-1 832/3 SNAP-GIPR cells expressing mini-G_s_-Venus or empty mVenus as a control in response to 100 nM GIP; *n*=3 independent time-lapse acquisitions. (**E**) Quantification of β2AR internalisation in Venus-positive *vs* −negative HEK293 SNAP-β2AR cells expressing mini-G_s_-Venus or empty mVenus as a control in response to 100 nM isoproterenol; *n*=3 independent time-lapse acquisitions. (**F**) Quantification of FFA2R internalisation in Venus-positive *vs* −negative HEK293 T-REx SNAP-FFA2R cells expressing mini-G_q_-Venus or empty mVenus as a control in response to 1 mM proprionate; *n*=3 independent time-lapse acquisitions. (**G**) Quantification of μ-OR internalisation in Venus-positive *vs* −negative HEK293 SNAP-μ-OR cells expressing mini-G_i_-Venus or empty mVenus as a control in response to 1 μM DAMGO; *n*=3 independent time-lapse acquisitions. Data is mean ± SEM; **p<0.01 by one-way ANOVA with Dunnett’s test; ns (non-significant).

## Discussion

In this report we have unveiled a previously unappreciated blocking effect of co-expressing mini-G proteins with their cognate GPCRs in the internalisation of these receptors in response to agonist stimulation. This effect, which is present across different GPCR classes and to different degrees with all mini-G protein subtypes tested, results in the inhibition of normal intracellular localisation of the receptor along the endocytic pathway and therefore hinders proper quantification of the degree of signalling elicited by these receptors from these intracellular locations.

Our results also show that β-arrestin-2 recruitment to the GLP-1R is completely blocked by the presence of mini-G_S_. This is an important finding, as it reveals that β-arrestin-2 recruitment requires prior dissociation of the Gαs subunit from the receptor. This might be somewhat expected as G proteins and β-arrestins are thought to engage the same inter-helical cavity on the intracellular side of the receptor (Carpenter et al., 2016; Zhou et al., 2017), so that the binding of one precludes binding of the other (Gurevich and Gurevich, 2019). However, this observation argues against the possibility of concomitant G protein- and β-arrestin-interacting “super-complexes”, which have been previously identified for other GPCRs (Thomsen et al., 2016), in the case of the GLP-1R when stimulated by its endogenous ligand.

Whilst the recruitment of β-arrestin-2 to the GLP-1R is essentially stopped in the presence of mini-G_s_, the same is not observed for GRK2: we were only able to detect a non-significant trend for a reduction in its recruitment to the receptor, suggesting that the stabilisation of the GLP-1R:mini-G_s_ complex affects GRK2 recruitment only partially at most. Additionally, there appears to be minimal differences in the segregation of GLP-1R to lipid nanodomains with and without mini-G_s_ co-expression. An interesting observation to note is that the basal level of GLP-1R found in lipid rafts is significantly greater with mini-G_s_ co-expression compared to mVenus controls, suggesting that the GLP-1R may already partially associate to lipid nanodomains in the absence of agonist stimulation when mini-G_s_ is co-expressed. Additionally, we could also observe recruitment of mini-G_s_ to AP2 hotspots at the plasma membrane following GLP-1 stimulation. Taken as a whole, these results demonstrate that GLP-1R nanodomain partitioning or interaction with AP2 is not dependent on β-arrestin-2 recruitment to the receptor, with these events instead likely potentiated by Gαs binding to active receptors.

We have also observed here that, at least for mini-G_s_, co-expression of the full-length G protein does not lead to the same blockage on GPCR internalisation as for its mini-G counterpart. While acute GPCR activation may occur in a similar manner with both, GPCR endocytosis appears to be prevented only with the latter. The most plausible explanation behind the β-arrestin-2 recruitment and receptor internalisation defects caused specifically by engineered mini-G_s_ proteins is the increased stability of the GPCR:mini-G interaction when compared to full-length G proteins (Carpenter and Tate, 2016). In normal conditions, binding of G proteins to cognate GPCRs results in the opening of the G protein nucleotide-binding pocket (Mahoney and Sunahara, 2016; Oldham and Hamm, 2008), leading to the loss of GDP in favour of the more abundant GTP (Gurevich and Gurevich, 2019). GTP-bound Gα proteins then dissociate from the receptor and from the Gβγ subunit, with both subunits then binding to their respective effectors. In the case of the mini-G proteins, however, the α5 helix at the C-terminus contains a mutation to improve the stability of the GPCR:mini-G complex (Wan et al., 2018). Additionally, movement of the α5 helix can rearrange the β6-α5 loop through modified α5-α1 interaction in the G protein (Mahoney and Sunahara, 2016). It has previously been observed that mutations that affect the β6-α5 loop can lead to the accelerated release of GDP, therefore increasing the GDP:GTP exchange rate of full-length Gαs (Iiri et al., 1994). It is therefore possible that the GPCR complex-stabilising α5 mutation present in mini-G_s_ proteins prevents GPCR deactivation through potentiating the release of GDP. While GRK recruitment may still occur almost normally under these conditions, prolonged GDP release could affect the degree of GRK-induced GPCR phosphorylation or simply hinder mini-G_S_ dissociation from the receptor despite near-normal receptor phosphorylation levels. This could in turn lead to β-arrestin recruitment blockage and significantly halt GPCR internalisation. Further studies looking into the G protein-coupling molecular mechanism could help elucidate this.

Mutations in the full-length Gαs β6-α5 loop also lead to constitutively activated adenylyl cyclase *in vitro*, and “hyperactive G protein signalling” associated with alterations in endocrine function (Iiri et al., 1994). While we have not observed significant differences in acute cAMP generation in mini-G_S_-expressing cells, perhaps due to increased signal amplification in our artificial receptor-overexpressing cellular systems, the observed blockage in desensitisation, and therefore also possibly resensitisation, of the GLP-1R in the presence of mini-G_s_ proteins could in turn potentially lead to prolonged activity, affecting cellular processes due to continuous activation of the GPCR in the absence of receptor internalisation and subsequent downregulation. The long-term consequences of this phenomenon remain to be established.

While the endocytosis of GPCRs that couple preferentially to mini-G_i_ and mini-G_q_ protein subtypes is also affected by the co-expression of their cognate mini-G proteins, the degree of this disruption appears to be dependent on the specific receptor tested. The reasons for this remain unclear and need to be further explored in detail, although might be related to the degree of stabilisation afforded to the specific GPCR:mini-G complex.

Despite differences in the overall effects observed between different receptors, an unexpected outcome of this investigation is the discovery of a close correlation between the coupling to each G protein subtype and the degree of GPCR internalisation inhibition elicited by co-expression of the corresponding mini-G. This could be exploited in the future to rapidly and cheaply assess the degree of coupling of any GPCR to different G proteins in any cell system amenable to transfection or viral transduction by simply co-expressing a tagged GPCR with a range of mini-G-Venus subtypes and assessing receptor internalisation rates in Venus-positive *versus* −negative cells for the GPCR of interest, avoiding the use of NanoBRET or NanoBiT assays which involve precise titration of the assay components and the use of expensive bioluminescent substrates.

Our results have important consequences for the interpretation of data on GPCR subcellular signalling when using a mini-G-based approach. As mini-G proteins are being used to detect the presence of activated GPCRs at different subcellular compartments, with the interrogation of endocytic signalling being a common application (Crilly et al., 2021; Jimenez-Vargas et al., 2020; Wan et al., 2018), there is clear potential for any mini-G-mediated alteration of receptor trafficking responses to confound these measurements. In view of the present observations, we recommend exhaustive checks on specific receptor internalisation rates if a mini-G is to be utilised in any intracellular signalling assay, and to consider changing to other non-mini-G based bystander methods including the use of nanobodies against G protein:active GPCR pairs, amongst other possibilities, such as, for example, the use of full-length G protein fusion constructs (Novikoff et al., 2021), or strategies to inhibit GPCR internalisation (Sposini et al., 2017).

## Supplementary Figure Legends

**Supplementary Figure 1: Lack of effect of full-length Gαs protein co-expression on GLP-1R internalisation and of mini-Gs co-expression on GLP-1R binding affinity to GLP-1 and cAMP generation.** (**A**) Percentage of GLP-1R internalisation in YFP-positive *vs* −negative INS-1 832/3 SNAP-GLP-1R cells co-expressing full-length Gαs-YFP in response to 100 nM GLP-1; *n*=3 independent time-lapse acquisitions. (**B**) AUC quantifications from (A). (**C**) Total cAMP GLP-1 dose response curves measured by HTRF in HEK293 T-REx SNAP_f_-GLP-1R-SmBiT cells co-transfected with mini-G_s_-Venus or mVenus as a control; *n*=3. Data is mean ± SEM; ns (non-significant) by paired t-test.

**Supplementary Figure 2: Lack of effect of mini-Gs co-expression on lipid nanodomain cAMP generation and recruitment to plasma membrane AP2 hotspots.** (**A**) Total cAMP data (dose response curves, logEC50 and Emax comparisons) measured by HTRF in cell samples from Figure 4F; *n*=4. (**B**) TIRFM analysis of mini-G_s_-Venus *vs* mVenus localisation to AP2-HA plasma membrane hotspots in HEK293 T-REx SNAP_f_-GLP-1R cells following 2 min stimulation with 100 nM GLP-1. Data is mean ± SEM; ns (non-significant) by paired t-test.

**Supplementary Figure 3: Lack of effect of mini-G co-expression on EGFR internalisation.** Quantification of EGFR internalisation in Venus-positive INS-1 832/3 SNAP_f_-EGFR cells co-expressing either mini-G_s_- or mini-G_i_-Venus, or empty mVenus as a control in response to 100 μg/mL EGF; *n*=3 independent time-lapse acquisitions. Data is mean ± SEM; ns (non-significant) by one-way ANOVA with Dunnett’s test.

## Acknowledgements

We thank the Facility for Imaging by Light Microscopy (FILM) at Imperial College London for help with microscopy data analysis in Fiji, Dr. Johannes Broichhagen (Leibniz-FMP Berlin) for the SNAP-μ-OR plasmid and Dr Ivan Corrêa Jr (New England Biolabs) for his supply of cleavable SNAP-tag probes. A.T. is supported by grants from the MRC (MR/R010676/1), Diabetes UK and the European Federation for the Study of Diabetes, as well as support from the Commonwealth and the Lilly Research Award Program (LRAP) scheme. B.J. is supported by the MRC (MR/R010676/1), IPPRF scheme, European Federation for the Study of Diabetes, Society for Endocrinology, British Society for Neuroendocrinology, Diabetes UK, the Lilly Research Award Program (LRAP) scheme and the UK National Institute for Health Research (NIHR) Imperial Biomedical Research Centre (BRC). M.M.S. and E.W.T. thank BBSRC for support (grant number BB/S001565/1). A.I. was funded by the LEAP 20gm0010004 and the BINDS JP20am0101095 from the Japan Agency for Medical Research and Development (AMED), KAKENHI 21H04791 and 21H05113 from by the Japan Society for the Promotion of Science (JSPS), Daiichi Sankyo Foundation of Life Science, Takeda Science Foundation, The Uehara Memorial Foundation and the Tokyo Biochemical Research Foundation. The Section of Endocrinology and Investigative Medicine is funded by grants from the MRC, BBSRC, NIHR and is supported by the NIHR Biomedical Research Centre Funding Scheme. The views expressed are those of the author(s) and not necessarily those of the funder, the NHS, the NIHR or the Department of Health.

