## Supplementary figures and images for "Expression of mini-G proteins specifically halt cognate GPCR trafficking and intracellular signalling"

### Supplementary Figures

#### Supplementary Figure 1

A

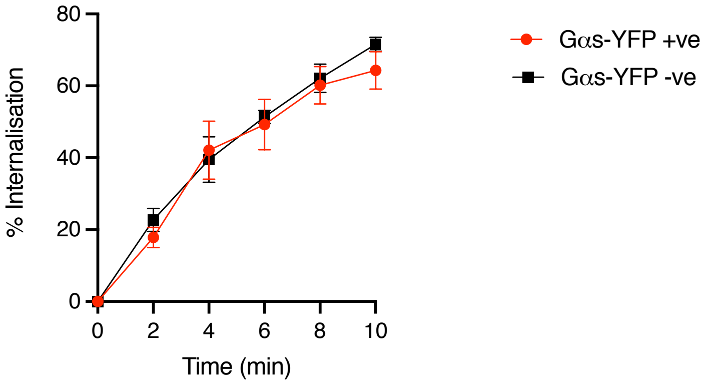


B

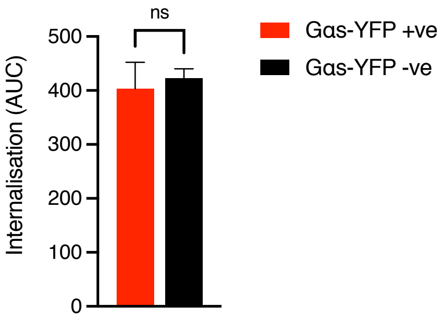


C

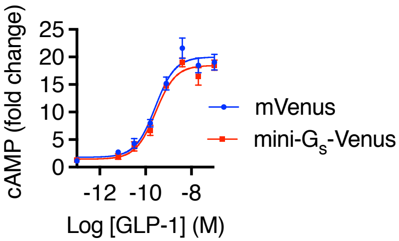

#### Supplementary Figure 2

**A**

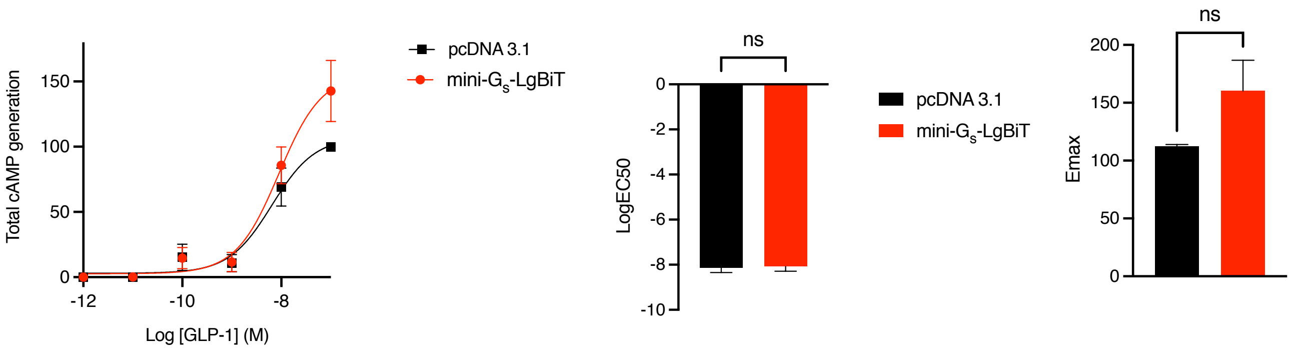


**B**

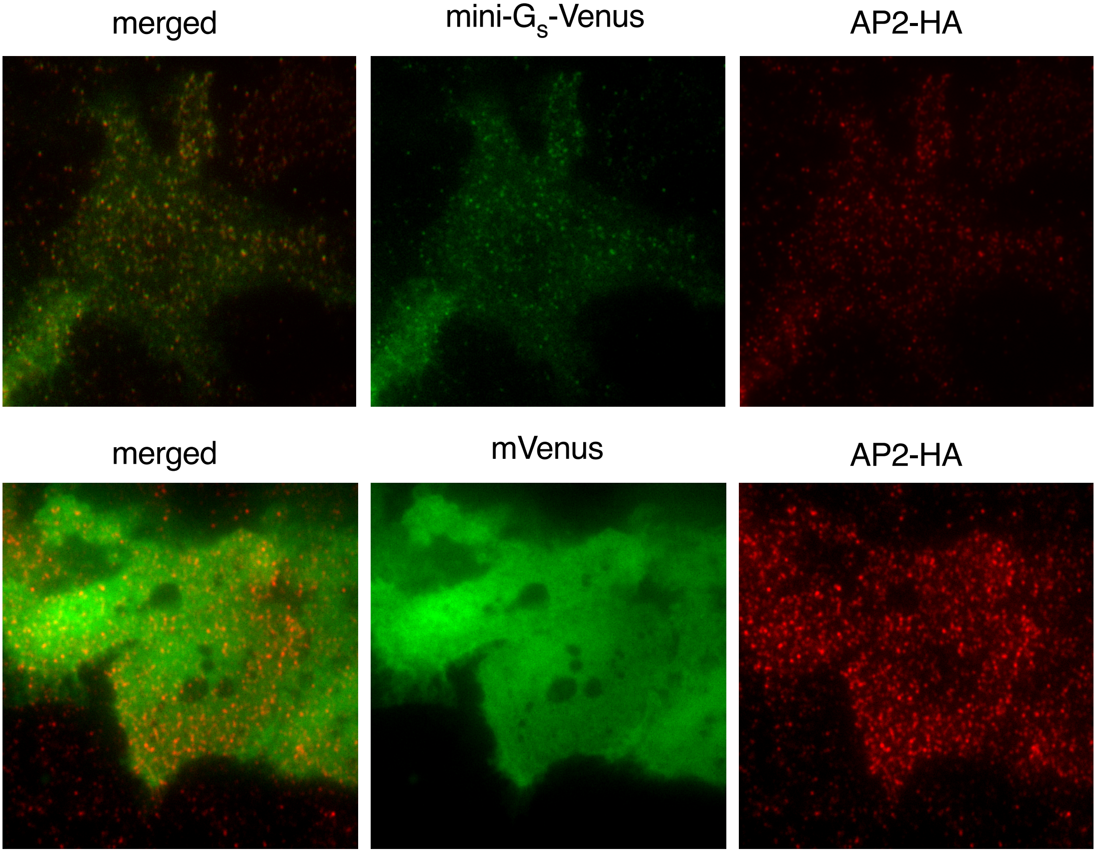

#### Supplementary Figure 3

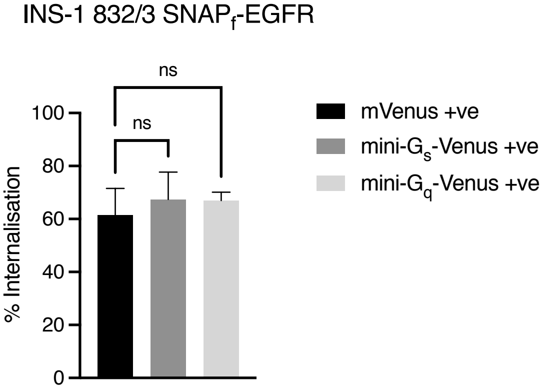
